# Hypoxia and epithelial to mesenchymal transition pathways are enriched in bladder tumor epithelium adjacent to tertiary lymphoid structures

**DOI:** 10.64898/2026.08.10.743535

**Authors:** Kartik Sachdeva, Priyanka Yolmo, Abdulhameed Abdulhamed, Gwenaëlle Conseil, Sadaf Rahimi, David M. Berman, Kathrin Tyryshkin, Roger Li, D. Robert Siemens, Madhuri Koti

## Abstract

Formation of tertiary lymphoid structures (TLS) within the bladder microenvironment because of chronic mucosal inflammation has been associated with variable clinical outcomes. While the immune cell composition and functional states of TLS have been characterized in both non-invasive and muscle-invasive bladder tumors, the TLS-adjacent tumor epithelial compartments remain poorly characterized. Evaluation of a 16-gene TLS signature in treatment-naïve tumor bulk RNA sequencing profiles from 283 non-muscle invasive bladder tumors, from patients treated with Bacillus Calmette-Guérin (BCG) immunotherapy, and 348 muscle-invasive bladder tumors from patients treated with immune checkpoint inhibitor therapy revealed overlapping enrichment of immune exhaustion pathways. High TLS gene expression scores correlated with upregulation of immune exhaustion, hypoxia, and epithelial-to-mesenchymal transition (EMT) pathways in tumors from both cohorts. Spatial whole transcriptomic analysis of tumor sections with high TLS density, revealed enrichment of genes associated with EMT, angiogenesis, extracellular matrix remodeling, and B cell receptor signaling pathways in tumor epithelial regions adjacent to TLS, whereas those distant from TLS exhibited enrichment of IFN-γ, TNF-α/NF-κB, p53, and metabolic pathways. Multiplex immunofluorescence further identified co-localization of exhausted immune cell populations within the core and periphery of peri-tumoral TLS. These findings indicate that a pro-tumorigenic microenvironment associated with disease progression in bladder cancer exists within peri-tumoral TLS and potentially a factor underlying contrasting therapeutic associations potentially driven by live microbial versus targeted immunomodulatory therapy in NMIBC and MIBC.

## Introduction

Bladder cancer (BC) is the ninth most frequently diagnosed urological malignancy worldwide and the most common cancer of the urinary tract[1–3]. Non-muscle invasive bladder cancer (NMIBC) represents the most prevalent form of bladder cancer, accounting for approximately 70-80% of all incident bladder cancer cases, whereas muscle-invasive bladder cancer (MIBC) comprises 20-30% of newly diagnosed BC[1, 2]. Bacillus Calmette-Guérin (BCG) immunotherapy remains the gold standard adjuvant treatment for NMIBC patients predicted to be at an increased risk of recurrence[4]. Several novel alternate therapies such as intravesical gemcitabine-docetaxel chemotherapy, programmed death-1 (PD-1) immune checkpoint inhibitor (ICI), adenoviral vectors, and oncolytic viruses, have been approved or are under advanced clinical trials for BCG unresponsive NMIBC[5–7]. For the treatment of MIBC, bladder-preserving trimodal therapy or radical cystectomy with neoadjuvant enfortumab vedotin and pembrolizumab are approved options for localized disease, while ICI targeting programmed cell death receptor-1(PD-1) or -its ligand 1 (PD-L1), enfortumab vedotin antibody-drug conjugate, and fibroblast growth factor receptor (FGFR)-targeted therapies are employed with palliative intent for advanced or metastatic disease[5, 8].

Therapeutic response in both NMIBC and MIBC is substantially governed by the pre-treatment immunological state of the tumor microenvironment. The tumor immune microenvironment (TIME) has thus been extensively studied towards biomarker discovery. Given the mucosal immune architecture and chronic inflammation, tertiary lymphoid structures (TLS) are a prominent feature of the bladder TIME and have emerged as promising prognostic and predictive biomarkers[9]. The composition of bladder mucosa-associated TLS varies with its maturation stages, similar to germinal centers within secondary lymphoid organs[10–12]. Their cellular functional states within the pre-treatment TIME reflect the broader host systemic immune physiology, shaped by chronic and persistent antigen exposure of microbial or non-microbial origins leading to the exhaustion of immune cells within TLS. Given that therapeutic success of intravesical BCG immunotherapy for NMIBC relies on recruitment of naïve immune cells and activation within the bladder microenvironment, pre-existing immune exhaustion limits its therapeutic efficacy[10, 12]. Our previous reports, using a retrospective cohort of patients with NMIBC and a carcinogen-induced aging model of bladder cancer, demonstrated an inverse association between high densities of exhausted tumor-infiltrating immune cells, B cell dominant TLS, and poor response to BCG[10, 12]. In contrast, higher TLS density and greater maturation state in MIBC tumors is associated with improved response to trimodal therapy and immune checkpoint inhibitor therapy[13–15].

The paradoxical associations of TLS in NMIBC and MIBC, presents a challenge in establishing their prognostic relevance biomarker potential[10, 12–14, 16, 17]. To bridge this gap, herein, we evaluated the prognostic relevance of a 16-gene TLS signature score using tumor transcriptome profiles from two large cohorts of patients with NMIBC and MIBC. To further delineate the spatial distinctions within transcriptional programs and pathways, we performed spatial whole transcriptomic profiling to map pathways enriched across epithelial regions topographically proximal and distal to TLS in high-grade NMIBC tumors. Multiplex immunofluorescence-based immunophenotyping was performed to spatially resolve the distribution and composition of innate and adaptive immune cell populations within the peri-tumoral TLS niche and adjacent tumor epithelium.

## Results

### 16-gene TLS signature-based stratification defines divergent therapeutic outcomes in NMIBC and MIBC

Patient stratification (n=283) based on TLS signature (*JCHAIN, IL7R, CD79A, FCRL5, MZB1, SSR4, XBP1, CD52, APOE, PTGDS, PIM2, DERL3, CXCL12, LUM, C1QA and C7*) scoring (**Fig. 1A**) was performed to interrogate its prognostic significance in NMIBC compared to MIBC. Kaplan-Meier survival analysis demonstrated that, within the BCG-treated NMIBC cohort[18], patients with high tumor TLS gene signature expression (n=141) scores exhibited significantly shorter high-grade recurrence-free (∼32% at 12 months) and progression-free survival (∼18% at 12 months) compared to those with low TLS gene signature scores (p < 0.05; **Fig. 1B, C**). The distribution of TLS scores across the patient cohort, arranged in ascending order, is illustrated in **Fig. 1D**.

**Figure 1.**
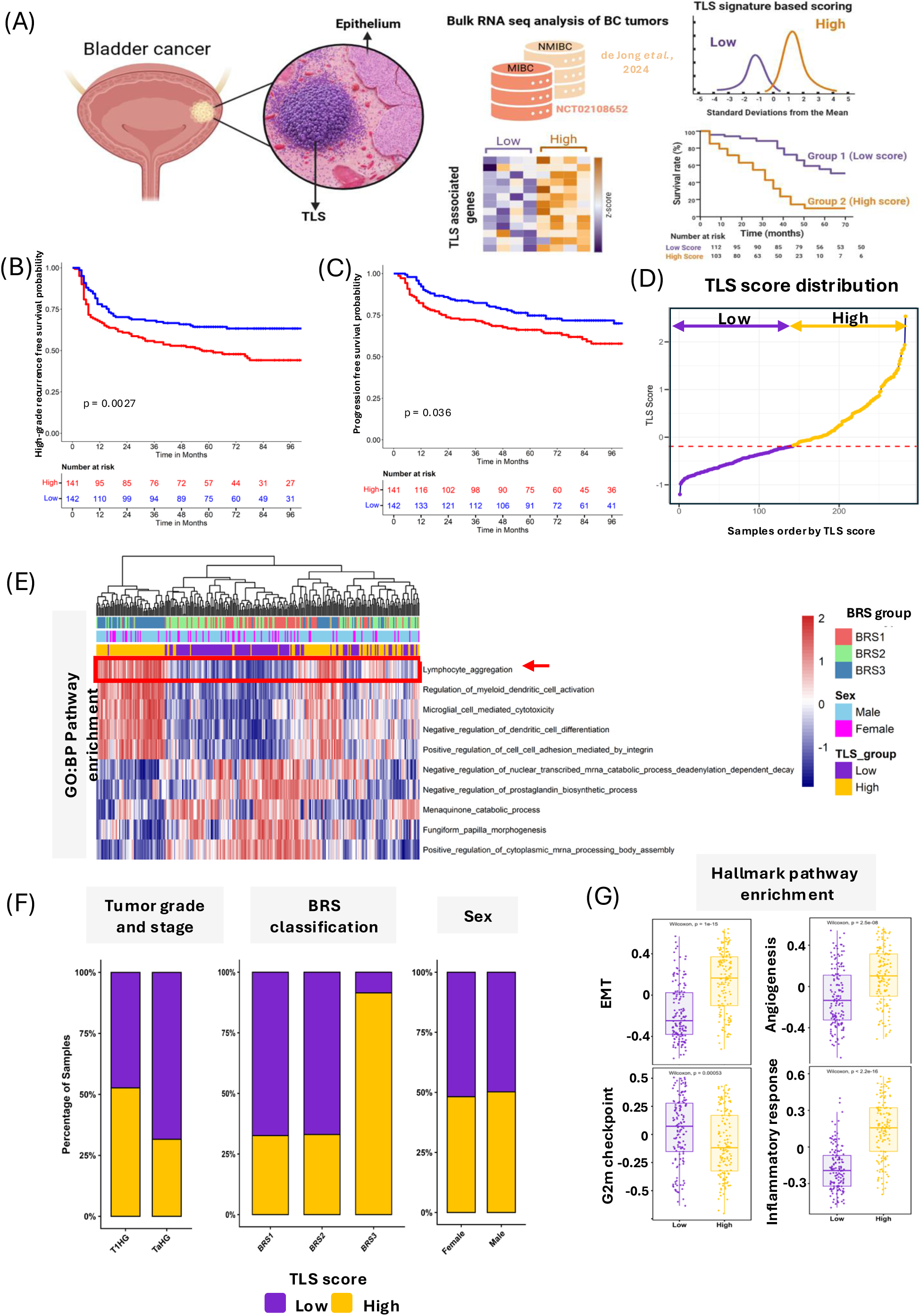
TLS gene signature score-based patient stratification in BCG-treated NMIBC and associations with clinical outcomes and biological pathways. Schematic overview of the analytical workflow employed to stratify patients with bladder cancer based on a 16-gene TLS signature score (A). Kaplan–Meier survival analyses demonstrating that patients with high TLS scores exhibit significantly greater risk of high-grade recurrence and earlier disease progression compared to those with low TLS scores in the BCG-treated NMIBC cohort (p < 0.05) (B, C). Line plot depicting the distribution of TLS scores across the patient cohort arranged in ascending order (D). Heatmap of GO:BP pathway enrichment Z-scores across TLS score groups, revealing preferential enrichment of immune-associated pathways including lymphocyte aggregation, negative regulation of dendritic cell differentiation, and integrin-mediated cell-cell adhesion in TLS-high tumors (E). Stacked bar plot illustrating the distribution of TLS score groups across clinical, biological, and transcriptomic characteristics, demonstrating predominance of high TLS scores in T1 high-grade tumors and the BRS3 molecular subtype, the latter associated with poor BCG therapeutic response (F). Hallmark pathway enrichment analysis demonstrating upregulation of EMT, angiogenesis, and inflammatory response pathways in TLS-high tumors, and enrichment of G2/M checkpoint signaling in TLS-low tumors, indicating divergent oncological transcriptional programs associated with TLS enrichment status (G).

To delineate the transcriptional basis underlying these clinical observations, pathway enrichment analyses were subsequently performed. GO:BP analysis identified significant enrichment of immune regulatory and lymphoid aggregation-associated pathways in TLS-high tumors (**Fig. 1E**; **Supplementary Table 2**), consistent with the establishment of an immunologically active yet potentially dysregulated tumor microenvironment. Furthermore, TLS scores were significantly elevated in T1 high-grade tumors relative to Ta high-grade tumors (**Fig. 1F**), corroborating the association between TLS enrichment and more advanced disease staging, in concordance with our previously published findings[19].

Molecular subtype analysis, based on the BCG response subtype (BRS) classification system[18], which stratifies NMIBC tumors based on transcriptomic profiles into three subtypes (BRS1, BRS2, and BRS3) associated with differential response to BCG immunotherapy, further revealed that approximately 85% of BRS3 tumors were classified as TLS-high, whereas BRS1 and BRS2 subtypes were predominantly represented within the TLS-low group (**Fig. 1F**). Hallmark pathway enrichment analysis demonstrated significant enrichment of EMT, angiogenesis, and inflammatory response pathways in TLS-high tumors (**Fig. 1G; Supplementary Table 2**). Conversely, TLS-low tumors exhibited preferential enrichment of cell cycle-associated pathways, most notably G2/M checkpoint signaling (**Fig. 1G; Supplementary Table 2**).

In the IMVigor210 cohort (locally advanced or metastatic urothelial cancer), TLS-high tumors also demonstrated elevated transcriptional expression of immune checkpoint genes, including *(PDCD1)* and *CD274* (**Fig. 2A**). In contrast to observations in the BCG-treated NMIBC cohort, patients with high TLS scores in the PD-L1 ICI treated IMVigor210 cohort exhibited a trend toward improved overall survival, though this association did not reach statistical significance (p<0.07) (**Fig. 2B**).

**Figure 2.**
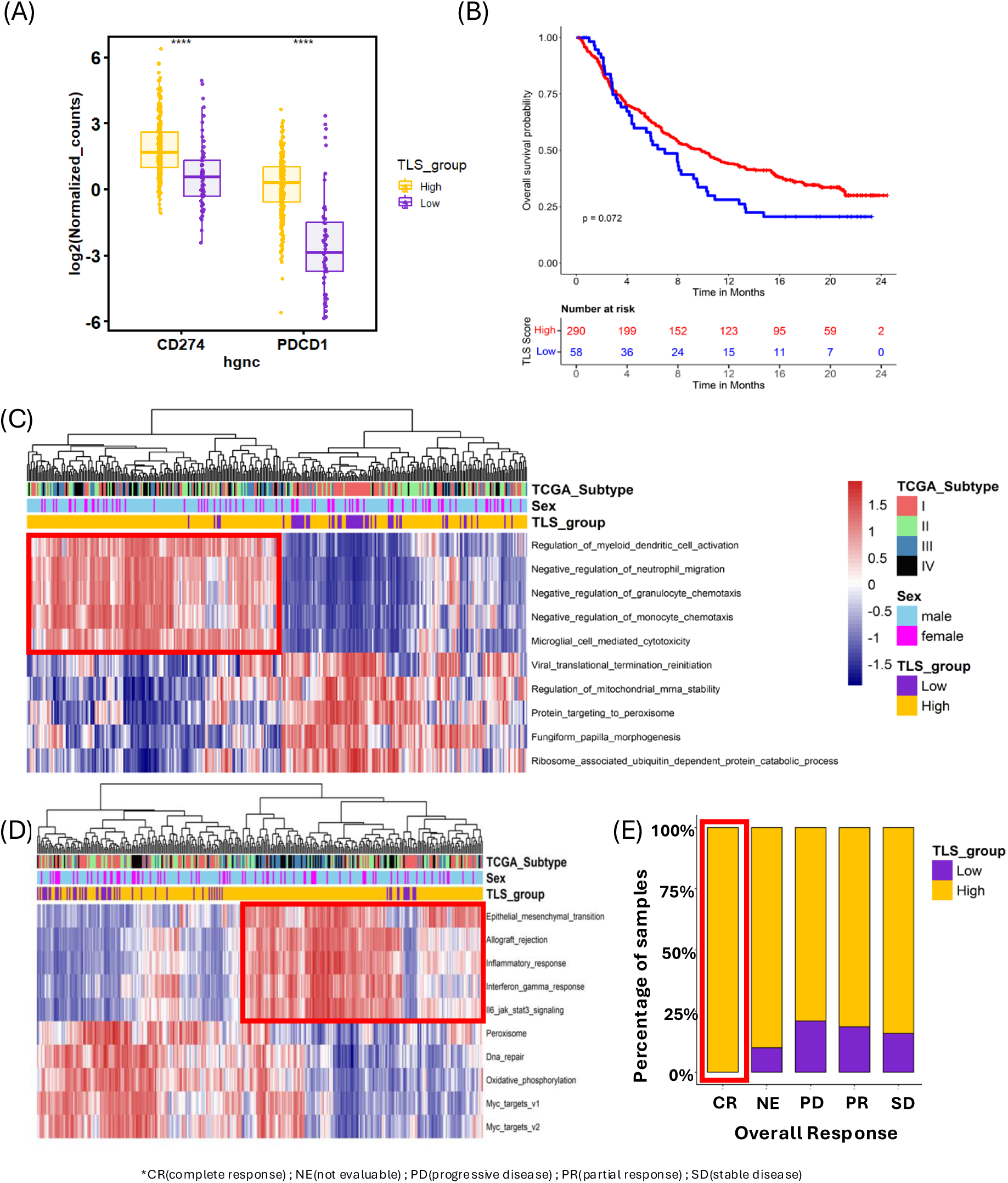
Validation of TLS gene signature scores in the atezolizumab-treated metastatic urothelial carcinoma cohort (IMVigor210). Boxplots depicting expression levels of *CD274* (PD-L1) and *PDCD1* (PD-1) across TLS score groups, demonstrating elevated immune checkpoint marker expression in the TLS-high patient subgroup, indicative of a pre-existing exhausted immune microenvironmental state (A). Kaplan–Meier survival analysis demonstrating a trend toward improved overall survival in patients with high TLS scores following atezolizumab treatment, suggesting a differential therapeutic benefit of immune checkpoint blockade in TLS-enriched tumors(B). Heatmap of GO:BP pathway enrichment z-scores across TLS score groups, revealing preferential enrichment of innate immune regulatory pathways in TLS-high tumors (C). Heatmap of Hallmark pathway enrichment z-scores, demonstrating upregulation of EMT, inflammatory response, interferon-γ signaling, and IL6–JAK–STAT3 pathways in the TLS-high patient group (D). Stacked bar plot illustrating the distribution of TLS score groups across clinical response categories, demonstrating that all patients achieving complete response to atezolizumab were classified within the TLS-high group, highlighting the potential predictive utility of TLS-based stratification in the context of immune checkpoint blockade (E).

Pathway enrichment analysis in the IMVigor210 cohort demonstrated upregulation of epithelial to mesenchymal transition (EMT), inflammatory response, interferon-γ signaling, and negative regulation of innate immune pathways in TLS-high tumors (**Fig. 2C, D; Supplementary Table 3**), similar to the transcriptional profiles observed in the BCG-treated cohort. Notably, all patients achieving a complete clinical response to atezolizumab were classified within the TLS-high group (**Fig. 2E**). These findings demonstrate that TLS-based patient stratification delineates biologically distinct tumor subgroups characterized by immune regulatory and exhaustion-associated transcriptional programs (**Fig. 1E–G**; **Fig. 2C–D**), with divergent prognostic and therapeutic implications associated with live microbial versus antibody-based immunomodulatory therapy.

Since older age significantly associated with progression free survival in the de Jong cohort we investigated whether TLS signature score also aligns with patient age. Our analysis of TLS score across the patient cohort did not reveal a significant relationship with older age. Stratifying samples into three age groups (≤50 years, 51–65 years, and >65 years) showed no statistically significant differences in TLS score distribution (Kruskal-Wallis, p= 0.49), and direct correlation analysis confirmed the absence of a linear association between age and TLS score (R = 0.0038, p = 0.95) (**Supplementary Figure 1**). Similarly, the proportion of samples classified as TLS-high versus TLS-low was broadly comparable across age groups, without a clear age-associated trend. These findings suggest that, within this cohort of high-risk NMIBC, TLS formation and abundance are not driven by chronological age.

### Epithelial to mesenchymal transition, hypoxia and angiogenesis pathways are enriched within epithelium adjacent to TLS

To further investigate the contrasting clinical associations of TLS enrichment in NMIBC, spatial transcriptomic profiles of tumor epithelial regions proximal to TLS were compared to epithelium with distant or no TLS across 10 whole-tumor sections from patients who experienced recurrence (**Fig. 3A, B**). GO:BP analysis further supported these observations, identifying enrichment of B cell receptor signaling, extracellular matrix organization, and extracellular structure remodeling pathways in TLS-adjacent epithelium (**Fig. 3C**). In contrast, the TLS-distant epithelial regions were characterized by enrichment of primary metabolic processes, cellular organization, translational regulation, and ribosome biogenesis (**Fig. 3C**), reflecting a metabolically active but relatively immune-excluded transcriptional state. Hallmark pathway analysis revealed that TLS-adjacent epithelium was enriched in hypoxia, EMT, and angiogenesis pathways **(Fig 3D)**. In contrast, TLS-distal epithelial regions were marked by interferon-γ response, TNF-α signaling via NF-κB, p53 pathway activation, and luminal marker expression, pointing to fundamentally divergent transcriptional states shaped by spatial proximity to TLS **(Fig 3D)**.

**Figure 3.**
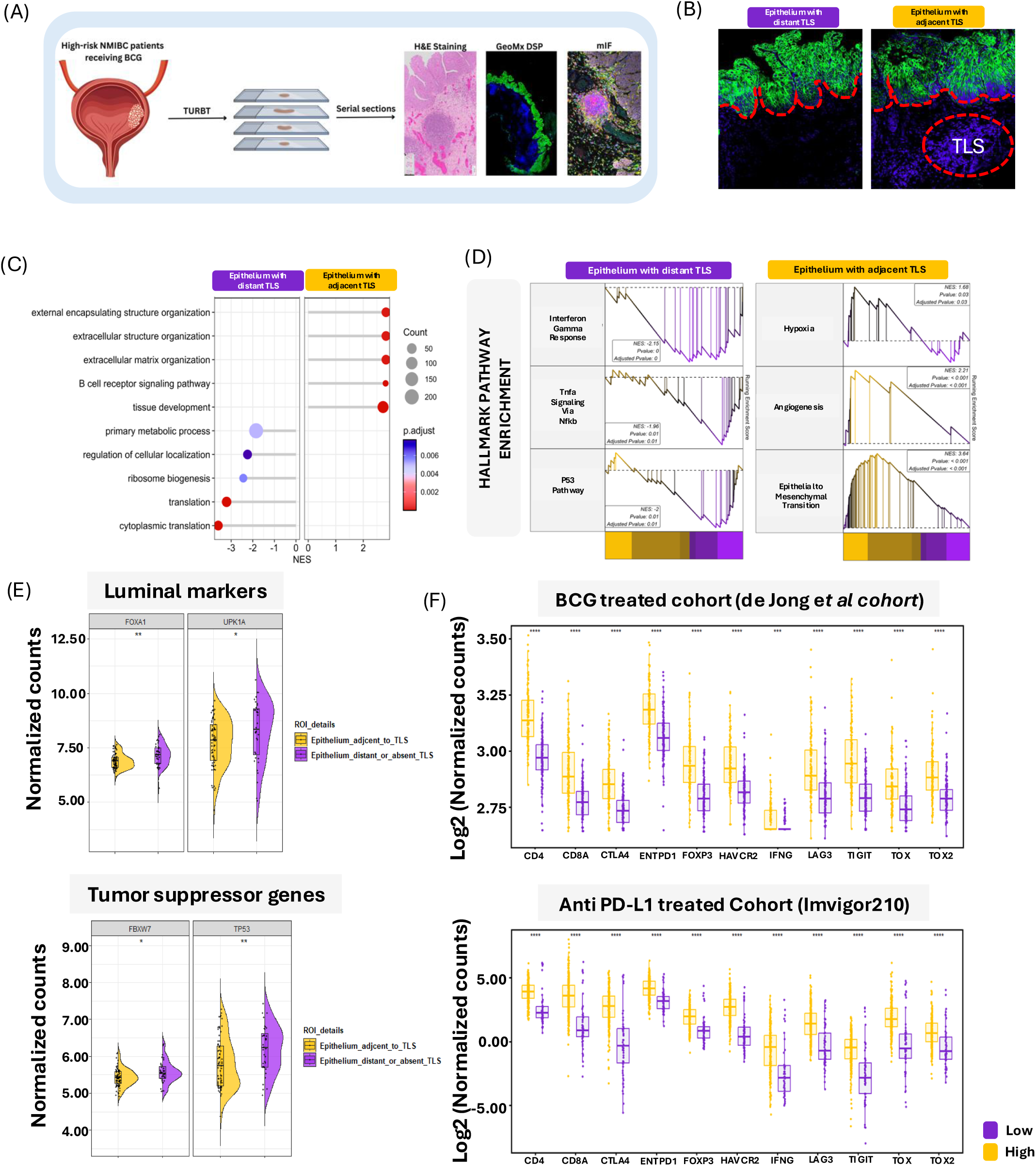
TLS-adjacent epithelium exhibits remodeling and immune-associated transcriptional programs in high-grade bladder cancer. Representative spatial transcriptomic images illustrating ROI selection across 10 whole-tumor sections, delineating tumor epithelial regions proximal to TLS and those distal or lacking adjacent TLS for comparative transcriptomic analysis (A, B). Hallmark and GO:BP pathway enrichment analysis comparing epithelial regions with adjacent and distant TLS, demonstrating preferential enrichment of hypoxia, EMT, angiogenesis, B cell receptor signaling, and extracellular matrix remodeling pathways in epithelium adjacent to TLS, and enrichment of interferon-γ response, TNF-α/NF-κB signaling, p53 pathway activation, and metabolic and translational programs in epithelium with distant or no TLS (C, D). Expression analysis of luminal differentiation markers *FOXA1* and *UPK1A* alongside tumor suppressor genes *TP53* and *FBXW7*, demonstrating significant downregulation in epithelium with adjacent TLS relative to epithelial regions with distant TLS, indicative of transcriptional alterations within epithelial regions in areas of active immune engagement (E). Validation across two independent bulk RNA sequencing cohorts demonstrating consistent upregulation of immune cell exhaustion-associated transcripts within the TLS-high patient group, reinforcing the transcriptional relevance of TLS-associated epithelial reprogramming at the population level (F).

Expression of *FOXA1* and *UPK1A*, established transcriptional markers of the luminal bladder cancer subtype, was significantly downregulated in TLS-proximal epithelial regions along with downregulation of tumor suppressor genes such as *TP53* and *FBXW7* (**Fig. 3E**), suggesting transcriptional differentiation away from the luminal phenotype in areas of active immune engagement. Immune cell exhaustion-associated transcripts were consistently upregulated within the TLS-high patient group in tumors from both the cohorts (**Fig. 3F**), reinforcing the broader transcriptional relevance of TLS-associated epithelial reprogramming beyond the spatial context. These findings suggest that TLS proximity may impose a distinct transcriptional identity on the adjacent tumor epithelium. Such a transcriptional state contrasts with the homeostatic and metabolic programs observed in TLS-distal epithelial regions.

### Spatial immunophenotyping reveals exhausted immune niches within peri-tumoral TLS

Spatial profiles of TLS intrinsic immune cell subsets were characterized via mIF across ROIs within the whole tumor sections (**Fig. 4A, B**). Examination of cell type distribution across the three spatially defined tissue compartments such as TLS-adjacent epithelium, TLS-distant epithelium, and lamina propria/tumor stroma, revealed compartment-specific enrichment of distinct immune populations (**Fig. 4C, D**). TLS-proximal regions were characterized by a higher proportion of B cells and PD1+ and CD4+FoxP3+ T cell populations, whereas TLS-distal epithelial regions and stromal compartments exhibited divergent immune compositions.

**Figure 4.**
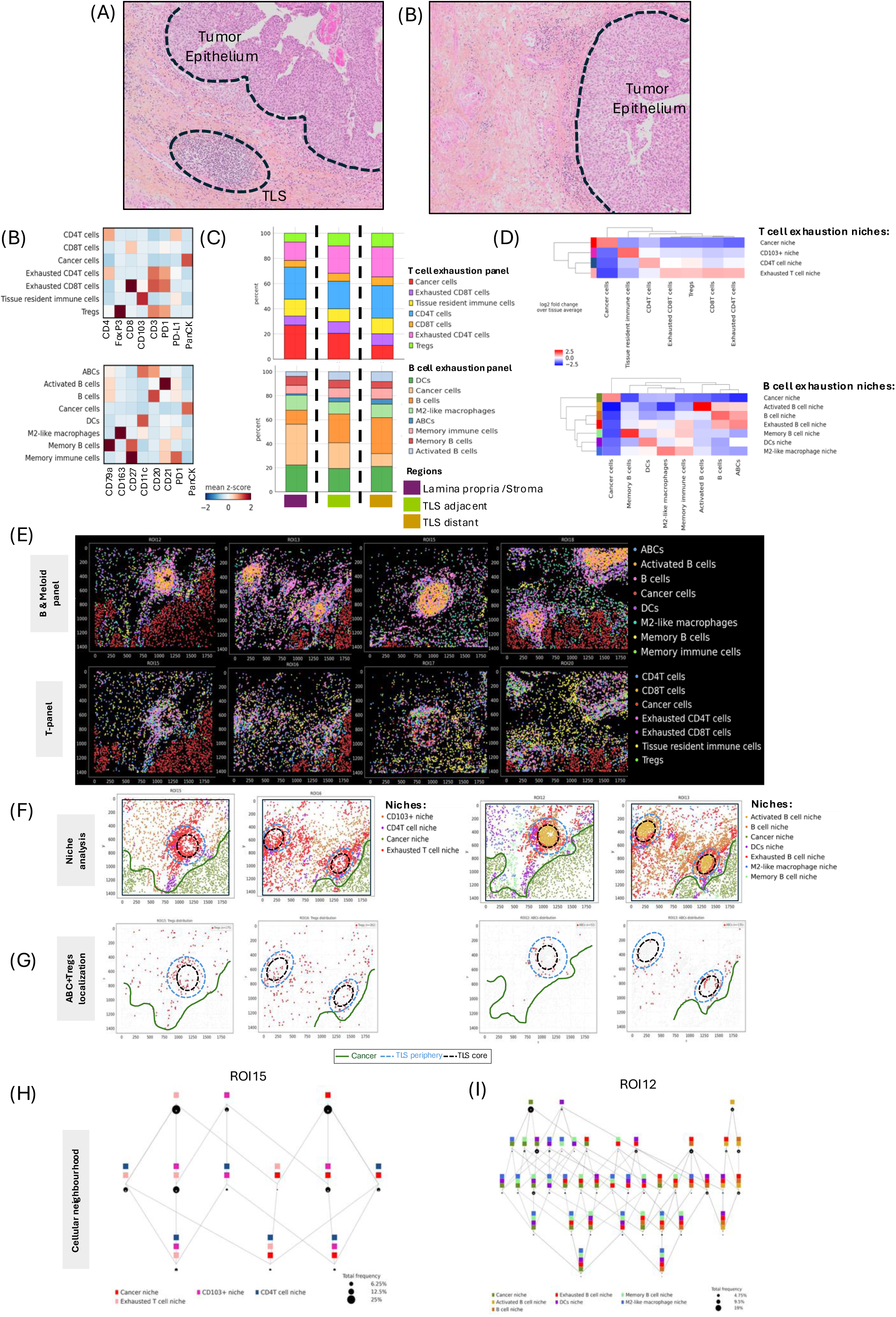
Spatial immunophenotyping of peritumoral TLS and adjacent tumor epithelium in high-risk bladder cancer tumors from non-responders. Representative H&E and corresponding mIF images illustrating morphological context and spatial distribution of selected ROIs across TLS and adjacent epithelial compartments (A, B). Marker expression with z-scores following spectral compensation for Panel 1 (B cell and myeloid) and Panel 2 (T cell), for cell phenotyping (C). Proportional distribution of annotated immune and epithelial cell populations across TLS-adjacent to epithelium, TLS-distant to epithelium, and other region (lamina propria/tumor-stroma compartments) (D). Heatmap depicting cell type enrichment within identified cellular neighborhood niches, defining discrete immune niches across both panels (E). Dot plots illustrate spatial distribution of annotated cell types across TLS and TLS-adjacent epithelial regions, reflecting differential immune composition between compartments (F). Dot plots depicting distribution of immune niches across TLS and adjacent epithelial regions, highlighting preferential enrichment of exhausted and B cell-associated niches in TLS-proximal compartments (G). Representative spatial scatter plots from ROI15 and ROI12 showing preferential localization of Tregs and ABCs at the TLS periphery relative to adjacent epithelial and distal regions (H). Spatial neighborhood analysis demonstrating minimal interaction between activated B cell and dendritic cell niches within TLS-rich regions, indicative of impaired antigen-driven B cell activation and functionally compromised humoral immunity within peritumoral TLS (I, J).

Unsupervised cellular neighborhood analysis identified discrete immune niches within the tumor microenvironment, with niche identities assigned based on the predominant cell type composition of each neighborhood (**Fig. 4E**). Within the T cell panel, four distinct niches were identified: CD103⁺; CD4⁺ T cell; cancer cells; and exhausted T cell niches. The B cell and myeloid panel resolved seven niches: activated B cell; total B cell; cancer cells; DCs; exhausted B cell; anti-inflammatory macrophage; and memory B cell niches. Dot plot visualization of cell type distributions across the four spatially defined TLS-associated regions following cell annotation further highlighted the differential enrichment of immune populations across TLS core, periphery and adjacent epithelium (**Fig. 4F**). Niche distribution analysis across these regions similarly demonstrated spatially restricted enrichment of specific immune niches, with exhausted and B cell-associated niches disproportionately represented in TLS-proximal compartments (**Fig. 4G**).

Regulatory T cells (Tregs) and exhausted-B cells such as atypical B cells (ABCs) exhibited preferential localization within the TLS periphery, as illustrated in representative ROI examples (**Fig. 4H**). This spatial enrichment of Tregs and ABCs at the TLS periphery is consistent with the establishment of a locally immunosuppressive microenvironment that may actively constrain effective anti-tumor immunity within peri-tumoral TLS. Representative cellular neighborhood analysis further illustrates the spatial organization and co-localization patterns of key immune cell populations. Activated B cell niches demonstrated minimal spatial interaction with DC niches within the peri-tumoral TLS microenvironment (**Fig. 4I, J**). In summary, spatial immunophenotyping revealed distinct, compartmentalized exhausted immune niches marked by peripheral enrichment of Tregs, ABCs and limited co-localization of activated B cells and DCs within peri-tumoral TLS.

## Discussion

The distribution and composition of TLS within the TIME represents a complex immune microenvironment associated with variable clinical outcomes in solid tumors including both NMIBC and MIBC. Aligning with the unique mucosal architecture of the bladder, increased TLS presence is known to associate with advancing disease stage and immune exhaustion[10, 12, 19–21]. While prior investigations have predominantly investigated TLS intrinsic immune cell composition, spatial architecture, and prognostic correlations[10, 13, 19, 22], the interactive phenotypes with the neighboring tumor epithelial compartment have received little attention. The present study addresses this critical gap in knowledge via characterizing the transcriptional states of tumor epithelium adjacent to and distant from TLS in tumors from BCG unresponsive NMIBC. The spatial convergence of pathways such as EMT, angiogenesis, hypoxia, extracellular matrix (ECM) remodeling, and immune exhaustion transcriptional programs in epithelium with adjacent TLS collectively delineates a topographically restricted pro-tumorigenic niche whose molecular characteristics may underlie cancer progression and downstream unfavorable clinical outcomes associated with elevated TLS gene signature scores particularly in BCG-treated patients with NMIBC.

Our prior investigations established that TLS in bladder cancer is associated with higher tumor grade and pathological stage[19]. Tumor bulk transcriptomic analysis of tumors from the BCG-naïve de Jong cohort[18] further revealed that tumors within BRS3 were enriched for B cell associated and immune exhaustion associated genes indicating that immune infiltration within this subtype does not necessarily culminate in functional antitumor activity following BCG therapy[10]. Further, systemic and tumor-intrinsic expansion of ABCs within TLS indicated local and systemic humoral immune dysfunction in BCG treated high-risk NMIBC[12]. An interesting observation in our study was the lack of association between older age and TLS-signature score. Older age is a known independent predictor of response to BCG as previously reported in the de Jong cohort and by others due to aging associated immune dysfunction and dampened response to vaccines and infection[23–28]. Our findings suggest chronic bladder mucosal inflammation associated TLS formation as another major contributor to poor response. Supporting these observations, we speculate that variable outcomes across different age groups and high TLS score could be driven by co-morbidities that promote chronic pre-BCG TLS formation and increased immune exhaustion in the bladder mucosa of patients <65 years of age.

Accumulation of Tregs within the tumor microenvironment is known to elevate local TGF-β levels and initiate EMT induction, establishing a mechanistic link between Treg infiltration and the acquisition of invasive capacity by adjacent epithelial cells. TGF-β drives EMT by disrupting cell-cell adhesion and activating core EMT effectors including SNAI1, ZEB1/2, and TWIST[29, 30]. In parallel, the tumor microenvironment promotes ECM remodeling through upregulation of collagen synthesis and matrix-modifying enzymes such as matrix metalloproteinases and lysyl oxidase family members, altering stromal architecture and tissue stiffness[31]. The spatial enrichment of ECM organization and extracellular structure remodeling pathways in TLS-adjacent epithelium identified in this study supports the interpretation that peri-tumoral immune aggregates, despite their morphological resemblance to organized lymphoid structures, paradoxically sustain a pro-invasive stromal niche within the bladder tumor microenvironment. Angiogenesis and hypoxia constitute further mechanisms through which the exhausted peri-tumoral immune niche reshapes both epithelial and stromal compartments. Within hypoxic intratumoral regions, HIF-1α activation is known to drive transcriptional upregulation of VEGF, inducing aberrant neovascularization characterized by structurally irregular and highly permeable vessels that impede effective immune cell trafficking into the tumor[32, 33]. Under hypoxic conditions, exhausted CD8⁺ T cells also adopt a pro-angiogenic secretory profile dominated by VEGF-A production, which in turn accelerates the differentiation of terminally exhausted T cell subsets, thereby exposing a reciprocal regulatory axis linking hypoxia, pathological angiogenesis, and immune exhaustion[34]. The spatial co-enrichment of hypoxia and angiogenesis pathway signatures in TLS-adjacent epithelium, therefore, correlates with a model in which peri-tumoral TLS resides within a hypoxic vascular niche where VEGF-mediated neovascularization, immune exhaustion, and epithelial remodeling reinforce one another within a unified pro-tumorigenic program.

B cell-dominant TLS have been linked to favorable clinical outcomes in MIBC, in contrast to the suppressive phenotype reported in BCG-treated NMIBC, highlighting a treatment context-dependent difference in their clinical association[14, 35, 36]. Stratification of the atezolizumab-treated IMVigor210 cohort by TLS gene signature score revealed a directional association between high TLS scores and improved overall survival, and all patients attaining complete clinical response were concentrated within the TLS-high stratum, reinforcing the therapeutic context-dependency of TLS-associated immune states. This is consistent with prior histology-based studies in MIBC treated with ICI: in a neoadjuvant durvalumab plus tremelimumab trial, higher baseline TLS density in pretreatment tumor specimens was associated with better response, longer relapse-free survival, and overall survival[16]. Furthermore, our findings are in concordance with a recent report where similar signatures and functional heterogeneity within TLS presence was shown to associate with treatment outcomes across various treatment modalities within chemotherapy treated and the IMVigor clinical trials[37].

Spatial immunophenotyping corroborated the immunoregulatory feature of TLS-rich peri-tumoral niches, revealing the predominance of CD163+ macrophages, antigen-experienced and exhausted B cell populations, and immunosuppressive and exhausted T cell subsets within these regions. The preferential spatial accumulation of regulatory T cells and ABCs at the TLS periphery, combined with the near-absence of DC-activated B cell spatial proximity within TLS-containing regions, points to a structural microenvironmental configuration. Given that DCs play a central role in TLS development and maintenance, the spatial uncoupling of DC and B cell populations within peri-tumoral TLS observed in our data may reflect a transitional or resolving stage of TLS maturation.

While novel findings from this study emphasize the significance of characterizing the epithelium adjacent to TLS in the context of the nature of immunomodulatory therapy, several limitations warrant acknowledgment. The spatial findings are derived from a numerically restricted single-institution cohort of BCG non-responders, and the cross-sectional study design precludes definitive causal attribution of epithelial remodeling to TLS-associated immune signals. Furthermore, the 16-gene TLS transcriptional surrogate may need further refinement to resolve the functional heterogeneity inherent to TLS of differing maturation states. Future investigations should prioritize validation in larger cohorts and mechanistic interrogation of TLS-epithelial crosstalk. Beyond stratifying between BCG and immune checkpoint inhibition, the rapidly expanding therapeutic landscape for BCG-unresponsive NMIBC including intravesical gene therapy (nadofaragene firadenovec), gemcitabine-docetaxel chemotherapy, IL-15 superagonist combinations (nogapendekin alfa inbakicept plus BCG) and oncolytic viral therapy (CG0070-cretostimogene grenadenorepvec) offer additional axes along which TLS gene signature-based stratification could be evaluated. Given that several of these agents act through IFNγ-driven or innate immune-activating mechanisms distinct from BCG’s reliance on adaptive effector recruitment, a pre-treatment TLS-signature may help identify patients more likely to benefit from these alternative immunumodulatory approaches.

## Materials and Methods

### Patient cohorts

This study was conducted in compliance with institutional ethical guidelines following approval by the Health Sciences Research Ethics Review Board at Kingston Health Sciences Centre (KHSC), Queen’s University. Publicly available bulk tumor transcriptomic profiles from two cohorts were accessed. The *de Jong cohort* was comprised of 283 treatment-naive tumors[18]. Normalized gene expression data for this cohort were obtained directly from the supplementary materials of the original publication[18]. The *IMvigor210* study included 348 patients with locally advanced or metastatic urothelial carcinoma treated with atezolizumab (programmed death ligand-1 antibody), representing a pooled subset of the original trial’s two clinical cohorts with available RNA sequencing data. Gene expression data for this cohort were accessed under EGA dataset accession EGAD00001003977[38]. For spatial whole transcriptomic profiling and immunophenotyping, whole tumor sections from 12 patients with high-grade bladder cancer who experienced recurrence following treatment with BCG at the KHSC were accessed (**Supplementary Table 1**).

### TLS signature score development and patient stratification

Normalized counts from publicly available RNA-sequencing profiles were used to evaluate the performance of TLS score-based patient stratification. A refined 16 gene signature (*JCHAIN, IL7R, CD79A, FCRL5, MZB1, SSR4, XBP1, CD52, APOE, PTGDS, PIM2, DERL3, CXCL12, LUM, C1QA and C7*) specific to TLS functional state was generated using the previously published comprehensive 29 gene set reflective of TLS composition and functional state evaluated in the *de Jong cohort[18, 39]* (**Supplementary Table 1**). Gene-level z-scores were calculated and averaged to derive a composite TLS score, with patients stratified into TLS-high and -low groups based on the median score. The expression data in both datasets were min-max standardized to correct for the differences. The Classification Learner App (MATLAB, v 2025a) was used to train all available machine learning classifiers to predict TLS status (high vs low) **Supplementary Table 2, Supplementary Figure 2**). The models were tested on a hold-out test set. Quadratic SVM (Model 2.12) was selected because it achieved consistent accuracy on the training and testing sets. The trained model was applied to the *IMvigor210 dataset* [40] to determine the TLS score per patient. The normalized expression data and associated clinical information for the *de Jong* NMIBC cohort were obtained directly from the original publication. For the *IMvigor210 MIBC cohort*, raw sequencing data and clinical information were accessed from the European Genome-Phenome Archive (EGA; https://ega-archive.org/). Sequencing reads from the *IMvigor210 dataset* were aligned to the human reference genome (GRCh38) using HISAT2 (v2.2.1). Gene-level quantification was performed using the featureCounts tool (Subread package, v2.0.6), generating raw count matrices. These counts were subsequently normalized to transcripts per million (TPM). TPM-normalized expression values were used for all downstream analyses.

### Pathway enrichment analysis, data visualization and statistical analysis

Gene set variation analysis (GSVA) was performed to estimate pathway-level enrichment across samples using the *GSVA R package* (v2.2.1). Hallmark (H) and gene ontology biological process (GO:BP) gene sets were obtained from the Molecular Signatures Database (MSigDB) using the *msigdbr package* (v25.1.1). Gene sets were converted into a list format, and GSVA scores were computed from normalized gene expression matrices using a Gaussian kernel function. Pathway-level associations with TLS scores were evaluated by correlating GSVA enrichment scores with TLS signature scores across samples (**Supplementary Table 2**).

To assess associations between TLS stratification and clinical or molecular variables, categorical distributions were visualized using 100% stacked bar plots, representing proportional differences across TLS groups. Additionally, boxplots were generated to compare TLS scores across clinical subgroups. For global pathway visualization, the top 10 most variable pathways were selected based on variance across samples. GSVA scores for these pathways were scaled (row-wise z-score normalization) and visualized using hierarchical clustering in heatmaps generated with the *pheatmap package* (v1.0.13). Sample annotations, including TLS group, sex, and molecular subtype, were incorporated to provide clinical context. All statistical analyses and visualizations were performed in R using standard packages including *ggplot2* (version 4.0.1) and *dplyr* (v1.1.4).

### Evaluation of clinical outcomes in NMIBC and MIBC using the 16-gene TLS signature score

Kaplan-Meier (KM) survival analysis was performed to determine associations of TLS score with recurrence free-, progression free- and overall survival. Survival time (in months) and event status were obtained from clinical metadata. Survival objects were constructed using the *survival package* (v3.8-3). Patients were stratified based on TLS score into TLS-high and -low groups, and survival distributions between groups were compared using the log-rank test. KM survival curves were generated using the *survminer package* (v0.5.1), with statistical significance assessed by two-sided p-values.

### NanoString GeoMx digital spatial whole transcriptomics profiling

To precisely determine the spatial whole transcriptomic profiles of TLSs and adjacent epithelial regions, serial 4 µm sections were prepared from archival Formalin-Fixed Paraffin-Embedded (FFPE) tumor specimens and mounted on charged slides at the Queen’s Laboratory of Molecular Pathology (QLMP). Unstained whole-tumor sections were processed on the NanoString GeoMx Digital Spatial Profiling (DSP) platform. Tissue sections were stained with a panel of fluorescently conjugated antibodies targeting pan-cytokeratin (PanCK) for tumor epithelial cell visualization, CD45 for immune cell identification, and SYTO13 for nuclear counterstaining. High-resolution fluorescence imaging was subsequently performed using the GeoMx DSP imaging system. Regions of interest (ROIs) were delineated based on PanCK immunoreactivity to demarcate tumor epithelial compartments, with consideration of spatial proximity to TLS. ROIs were manually annotated to encompass PanCK⁺ epithelial areas both proximal and distant to TLS, enabling comparative spatial transcriptomic analysis. A total of 170 ROIs were annotated across 10 tumor specimens, with a mean surface area of 182,893 µm² per ROI. Within each ROI, segmentation masks were applied to discriminate PanCK⁺ epithelial from PanCK⁻ non-epithelial compartments. Following quality control assessment and normalization using the NanoString GeoMx DSP software suite, 106 of the 170 ROIs were retained for downstream analysis, comprising 70 ROIs designated as TLS-proximal epithelium and 36 ROIs as TLS-distal epithelium. Transcriptomic data analysis was conducted within the R/Bioconductor environment using the standR package (v1.11.1), which facilitated a reproducible and standardized analytical workflow[41]. Raw count data alongside corresponding sample and feature annotations were imported into R and structured as a SpatialExperiment object via the readGeoMx() function. To minimize technical variability and ensure cross-sample comparability, Trimmed Mean of M-values (TMM) normalization was implemented using the geomxNorm() function.

Differential gene expression analysis between TLS-proximal and TLS-distal epithelial regions was conducted using the limma-voom pipeline, implemented within the StandR package framework for spatial transcriptomic data, following the GeoMx Analysis Workflow (Davis Laboratory; https://davislaboratory.github.io/GeoMXAnalysisWorkflow/articles/GeoMXAnalysisWorkflow.html). Log fold changes (logFC) between epithelial regions with and without adjacent TLS were computed using the same analytical approach. Genes meeting a minimum absolute logFC threshold of 0.25 and a nominal p-value ≤ 0.05 were retained for downstream gene set enrichment analysis (GSEA). GSEA was performed using the clusterProfiler package (v4.6.0) against Gene Ontology Biological Process (GO:BP) terms and Hallmark (H) gene sets from the Molecular Signatures Database (MSigDB) **(Supplementary Table 3)**. Enrichment scoring was computed using the fast gene set enrichment algorithm (fgsea), with gene set sizes constrained to a minimum of 3 and a maximum of 500 genes. Statistical significance was defined at a nominal p-value threshold of 0.05, applied without multiple testing correction given the exploratory nature of the spatial transcriptomic analysis. Enrichment results were visualized using the GseaVis package (v0.1.1; https://github.com/junjunlab/GseaVis).

### Spatial immunophenotyping using multiplex immunofluorescence assay

Spatial immunophenotyping was performed on 4 µm thick unstained FFPE sections using two distinct nine-plex antibody panels to interrogate B cell and myeloid exhaustion states (**Panel 1**), and T cell exhaustion states (**Panel 2**), respectively (**Supplementary Table 1**). Panel 1 comprised the following markers: CD20, CD79a, CD27, CD163, PD-1, CD11c, CD21, PanCK, and DAPI. Panel 2 comprised: CD8, CD3, FoxP3, CD103, PD-1, CD4, PD-L1, PanCK and DAPI (**Supplementary Fig. 3**). Both panels were applied using the Opal platform at the Molecular and Cellular Immunology Core Facility, Deeley Research Centre, BC Cancer Agency (Victoria, BC, Canada). Stained sections were acquired using the Vectra 3 multichannel imaging system (Akoya Biosciences), enabling high-resolution multichannel fluorescence image capture. Images were subsequently imported into Phenochart (v1.1.0; Akoya Biosciences) and QuPath (v0.4.2) for tissue annotation and high-magnification image capture. Positive staining thresholds for each marker were manually defined and confirmed using the DAPI nuclear counterstain channel.

Single-cell segmentation was performed using DeepCell Mesmer, a deep learning-based segmentation algorithm integrated within the SPACEc framework[42], which enables whole-cell and nuclear segmentation in highly multiplexed tissue images[43]. Fluorescence signal compensation was subsequently conducted using the compensation function implemented within SPACEc, which applies methodology derived from CellSeg[44]. Low-quality and non-cellular events were excluded based on predefined morphological and signal intensity filtering criteria to ensure downstream analytical integrity. A total of 167 ROIs were analyzed using Panel 1, and 166 ROIs were analyzed using Panel 2, across the full cohort.

Cell phenotype annotation was subsequently performed within SPACEc, whereby discrete immune and epithelial populations were classified based on marker co-expression profiles. Panel 1 enabled identification of B cell subpopulations, including total atypical B cells (ABCs; CD20+CD11c+CD21-), activated B-cells (CD79a+CD21+CD27-), cancer cells (PanCK+), memory B cells (CD79a+CD27+), as well as myeloid subsets including dendritic cells (DC; CD11c+) and immunosuppressive macrophages (CD163⁺). Panel 2 facilitated delineation of cytotoxic T cells (CD3⁺CD8⁺), helper T cells (CD3⁺CD4⁺), regulatory T cells (CD3⁺CD4⁺FoxP3⁺), and exhausted T cell subsets defined by PD-1 expression. PanCK positivity was used across both panels to demarcate tumor epithelial cells from immune populations. Following cell annotation, spatial neighborhood analysis was conducted to quantify immune cell composition, density, and co-localization patterns within TLS core, TLS periphery, and tumor epithelial compartments.

## Data availability

The data supporting the findings of this study can be found within the manuscript and its accompanying Supplementary files. Any additional materials or data are available from the corresponding author upon reasonable request.

## Author contributions

MK, RL and KS conceptualized and designed the study. KS, PY, and GC performed experiments included in this study, analyzed data, and contributed to manuscript writing and reviewing. KT guided transcriptomic data analysis and TLS signature development and validation. SR helped curate patient specimens. AA helped with reviewing histopathological features and retrieving archival tumor tissue specimens under the supervision of DMB. DRS and DMB helped with clinical classifications, study design, and contributed to patient recruitment and selection. All authors reviewed the manuscript.

## Acknowledgements

This study was supported by research operating grants from the Canadian Institutes of Health Research, Cancer Research Society, Leo and Anne Albert Institute for Bladder Cancer Care and Research. We are grateful to the Lab Research Services team in the Department of Pathology and Molecular Medicine at KHSC for their help in sectioning archival FFPE tumor tissue. We also thank QLMP for their support with slide scanning using Halo software, and Katy Milne of BC Cancer’s Molecular and Cellular Immunology Core (MCIC) for support with spatial immunophenotyping assays.

## Supplementary Figures

**Supplementary Figure 1.**
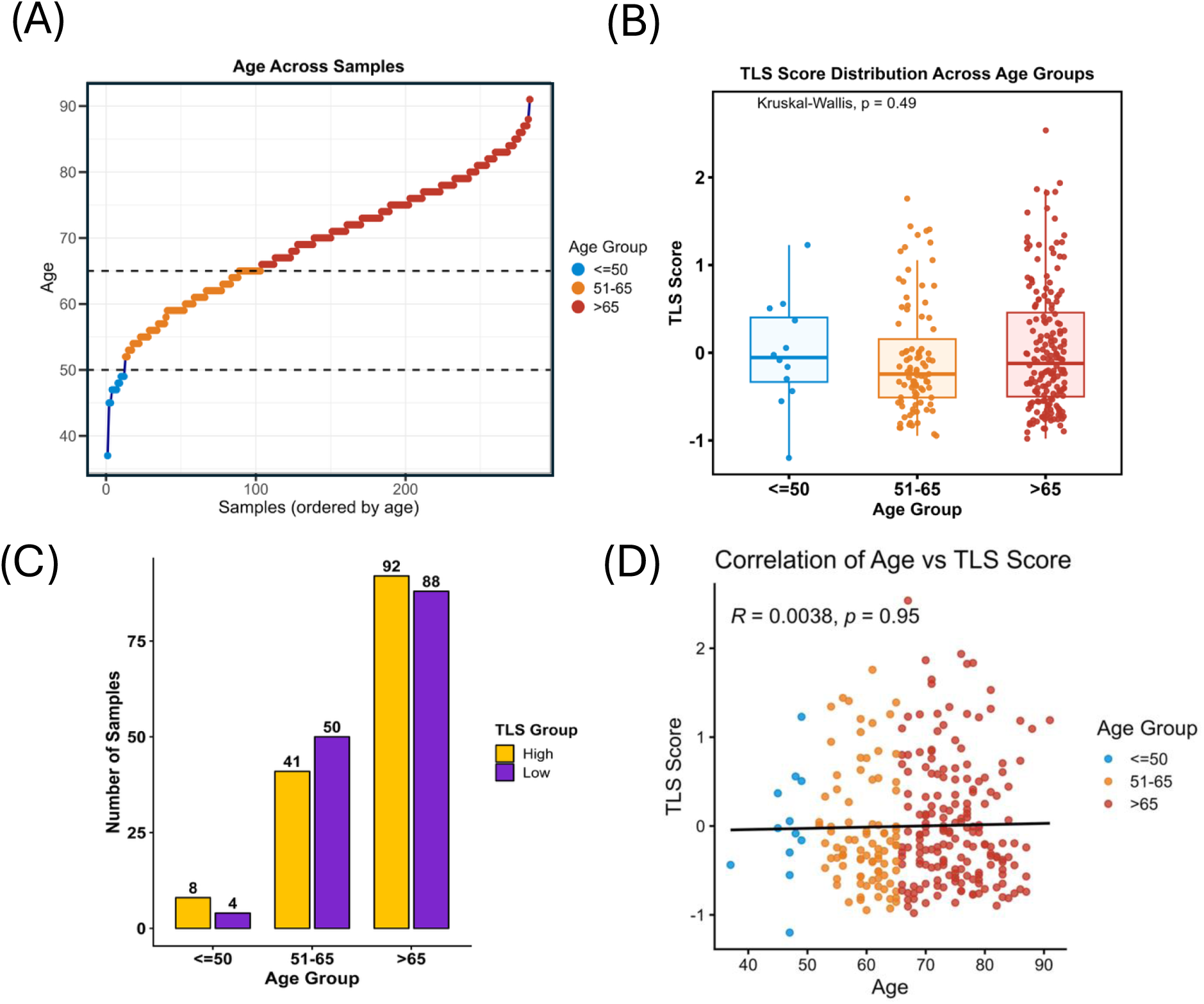
Relationship between patient age and TLS score. **(A)** Distribution of patient ages across the NMIBC cohort (de Jong *et al.,*), ranked in ascending order. Dashed horizontal lines mark the fixed age cutoffs (50 and 65 years) used to stratify samples into three age groups (≤50, 51–65, and >65, colored blue to red), with each point representing one sample. **(B)** Boxplots showing TLS score distributions within each age group. A Kruskal-Wallis test found no significant difference in TLS score across age groups (p value = 0.49). **(C)** Number of samples classified as having “High” (gold) or “Low” (purple) TLS scores within each age group. **(D)** Scatter plot of TLS score as a function of patient age, with points colored by age group. A linear regression line is shown in black. Pearson correlation analysis revealed no significant association between age and TLS score (R = 0.0038, p value = 0.95).

**Supplementary Figure 2.**
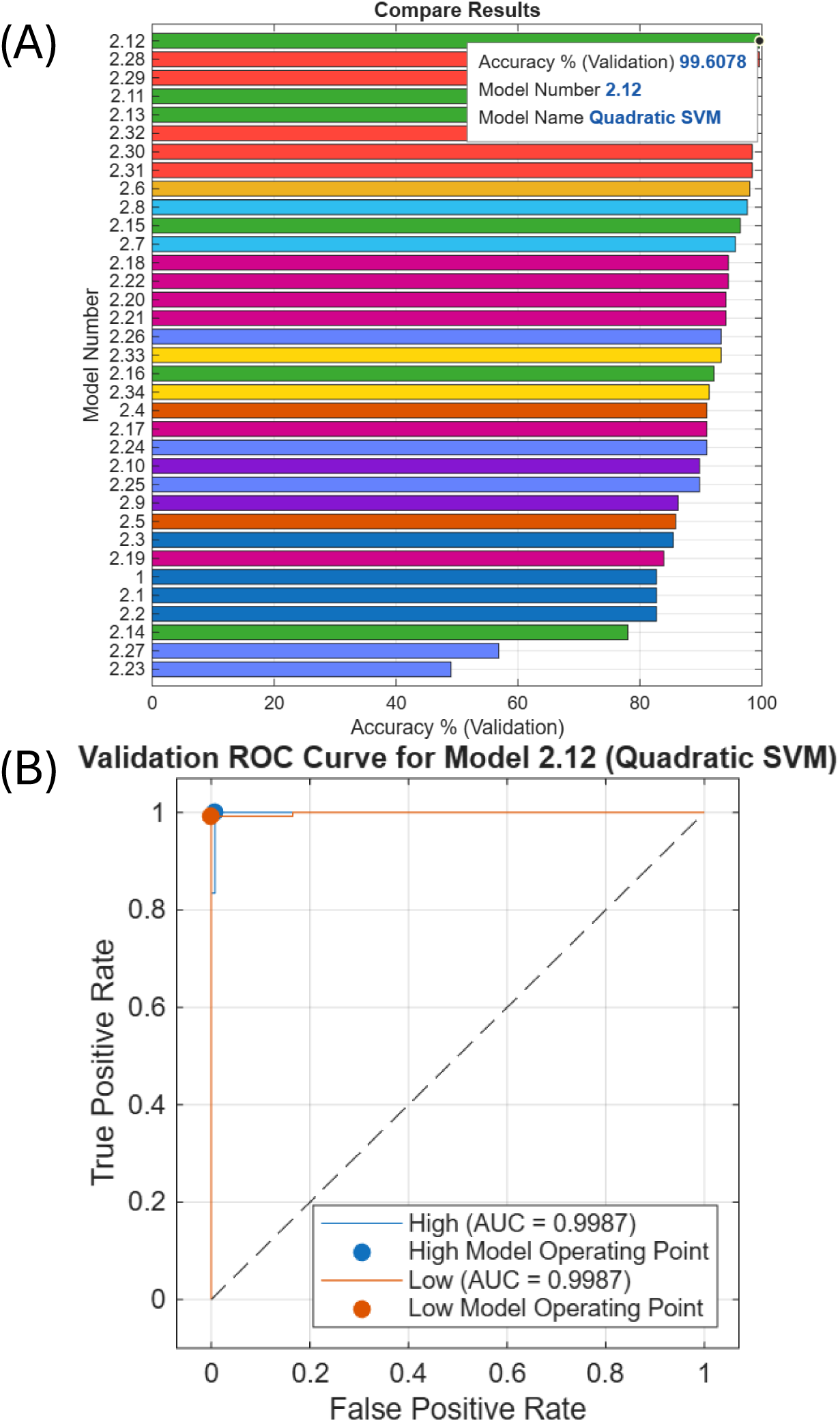
TLS score-based classifier. A) Model 2.12, a support vector machine (SVM) classifier, was identified as the best-performing model, achieving a classification accuracy exceeding 99%. (B) Receiver operating characteristic (ROC) curves for prediction of TLS-high and TLS-low groups, with an area under the curve (AUC) of 0.9987.

**Supplementary Figure 3.**
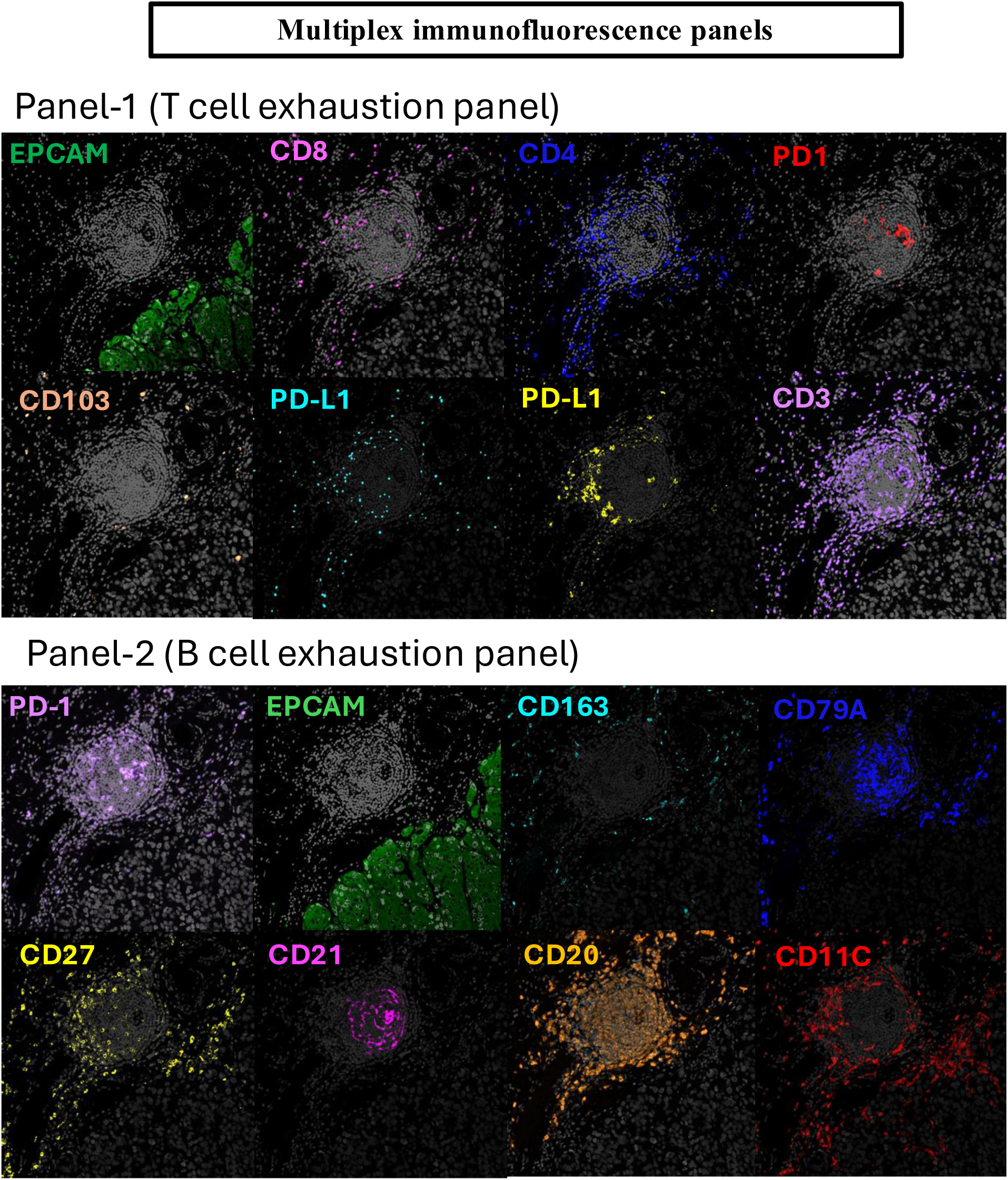
Localization of individual immune markers within tissue sections. Representative images showing spatial distribution and tissue localization of individual markers used in multiplexed immunofluorescence, highlighting distinct immune cell populations and their microanatomical context.

## Notes

https://ega-archive.org/dacs/EGAC00001000945

https://pmc.ncbi.nlm.nih.gov/articles/PMC10572776/#SM1

### Competing Interest Statement

The authors have declared no competing interest.

## References

1. Lenis, A., et al., Bladder Cancer—Review. JAMA, 2020. 324(19).

2. Dyrskjøt, L., et al., Bladder cancer. Nature reviews. Disease primers, 2023. 9(1).

3. Ferlay, J., et al. Global Cancer Observatory: Cancer Today. 2024.

4. van Rhijn, B., et al., Recurrence and progression of disease in non-muscle-invasive bladder cancer: from epidemiology to treatment strategy. European urology, 2009. 56(3).

5. St-Laurent, M., et al., Advances in the management of localized bladder cancers. Nature reviews. Clinical oncology, 2026. 23(3).

6. Jeong, S. and J. Ku, Treatment strategies for the Bacillus Calmette-Guérin–unresponsive non-muscle invasive bladder cancer. Investigative and Clinical Urology, 2023. 64(2).

7. Huang, X., et al., Mechanisms, Clinical Trials, and New Treatments for BCG-Unresponsive in Nonmuscle Invasive Bladder Cancer. Cancer Medicine, 2025. 14(18).

8. Cigliola, A., et al., The Management of Muscle Invasive Bladder Cancer: State of the Art and Future Perspectives. Cancers, 2025. 17(24).

9. van Dijk, N., et al., The Tumor Immune Landscape and Architecture of Tertiary Lymphoid Structures in Urothelial Cancer. Frontiers in Immunology, 2021. 12.

10. Yolmo, P., et al., Atypical B Cells Promote Cancer Progression and Poor Response to Bacillus Calmette-Guérin in Non–Muscle Invasive Bladder Cancer. Cancer Immunology Research, 2024. 12(10).

11. Chen, C., et al., B cells disrupt tertiary lymphoid structure formation and suppress anti-tumor immunity. Cancer Cell, 2026. 44(3).

12. Yolmo, P., et al., B cell exhaustion associates with poor response to Bacillus Calmette-Guérin immunotherapy in patients with bladder cancer. bioRxiv, 2026.

13. Hassan, N., et al., The Clinical Relevance of Tertiary Lymphoid Structures in Assessing Treatment Response in Muscle-Invasive Bladder Cancer. International Journal of Radiation Oncology, Biology, Physics, 2026. 124(3).

14. Teng, X., et al., Tertiary Lymphoid Structures as Independent Predictors of Favorable Prognosis in Muscle-Invasive Bladder Cancer. Cancer medicine, 2025. 14(10).

15. Zhou, L., et al., Tertiary lymphoid structure signatures are associated with survival and immunotherapy response in muscle-invasive bladder cancer. Oncoimmunology, 2021. 10(1).

16. Gao, J., et al., Neoadjuvant PD-L1 plus CTLA-4 blockade in patients with cisplatin-ineligible operable high-risk urothelial carcinoma. Nature medicine, 2020. 26(12).

17. Vidotto, T., et al., DNA damage repair gene mutations and their association with tumor immune regulatory gene expression in muscle invasive bladder cancer subtypes. Journal for Immunotherapy of Cancer, 2019. 7.

18. FC, d.J., et al., Non-muscle Invasive Bladder Cancer Molecular Subtypes Predict Differential Response to Intravesical Bacillus Calmette-Guérin. Science translational medicine, 2023. 15(697).

19. Koti, M., et al., Tertiary Lymphoid Structures Associate with Tumour Stage in Urothelial Bladder Cancer. Bladder Cancer, 2017. 3(4).

20. Pagliarulo, F., et al., Molecular, Immunological, and Clinical Features Associated With Lymphoid Neogenesis in Muscle Invasive Bladder Cancer. Frontiers in immunology, 2022. 12.

21. Grande, E., et al., Spatial architecture contributes to failure of bulk biomarker-guided neoadjuvant immunotherapy selection in bladder cancer: The DUTRENEO study. Cell Reports Medicine, 2026.

22. Ma, G., et al., Presence, Subtypes, and Prognostic Significance of Tertiary Lymphoid Structures in Urothelial Carcinoma of the Bladder. The Oncologist, 2023. 29(2).

23. Abou Chakra, M., et al., Impact of age on the effectiveness of intravesical Bacillus Calmette-Guérin and gemcitabine/docetaxel in treatment-naïve high-risk non-muscle-invasive bladder cancer. Translational andrology and urology, 2026. 15(6).

24. Inoue, T., et al., Association of Increased Age With Decreased Response to Intravesical Instillation of Bacille Calmette-Guérin in Patients With High-Risk Non-Muscle Invasive Bladder Cancer: Retrospective Multi-Institute Results From the Japanese Urological Oncology Research Group JUOG-UC-1901-BCG. Urology, 2022 Sep. 167.

25. Aghamir, S., et al., Oncologic outcomes of Bacillus Calmette-Guérin therapy in elderly patients with non-muscle-invasive bladder cancer: A meta-analysis. PloS one, 2022. 17(5).

26. Tavelli, J., et al., The impact of age on BCG treatment response. Urologic oncology, 2025. 43(7).

27. Koti, M., E. Michaud, and D. Siemens, Nonmodifiable Drivers of Response to Bacillus Calmette-Guérin: Considerations of Age, Sex, and Chronic Inflammation for the Management of Nonmuscle-Invasive Bladder Cancer. The Journal of urology, 2025. 214(4).

28. Poolman, J., Expanding the role of bacterial vaccines into life-course vaccination strategies and prevention of antimicrobial-resistant infections. NPJ vaccines, 2020. 5(1).

29. Hao, Y., D. Baker, and P. Ten Dijke, TGF-â-Mediated Epithelial-Mesenchymal Transition and Cancer Metastasis. International Journal of Molecular Sciences, 2019. 20(11).

30. Oh, E., J. Hong, and C. Yun, Regulatory T Cells Induce Metastasis by Activating Tgf-Â and Enhancing the Epithelial–Mesenchymal Transition. Cells, 2019. 8(11).

31. Yuan, Z., et al., Extracellular matrix remodeling in tumor progression and immune escape: from mechanisms to treatments. Molecular Cancer, 2023. 22(1).

32. Shi, S., et al., Research progress of HIF-1a on immunotherapy outcomes in immune vascular microenvironment. Frontiers in Immunology, 2025. 16.

33. Lee, W., et al., Combination of anti-angiogenic therapy and immune checkpoint blockade normalizes vascular-immune crosstalk to potentiate cancer immunity. Experimental & molecular medicine, 2020. 52(9).

34. Bannoud, N., et al., Hypoxia Supports Differentiation of Terminally Exhausted CD8 T Cells. Frontiers in Immunology, 2021. 12.

35. Penunuri, A., et al., The Emerging Role of B Cells and Tertiary Lymphoid Structures in Bladder Cancer. Current Urology Reports, 2025. 27(1).

36. Elias, R., et al., Intratumoral Expression of a Composite B Cell / CD8 T Cell Biomarker Stratifies Overall Survival by Circulating Tumor DNA Status and Benefit From Adjuvant Immunotherapy in High-Risk Muscle-Invasive Urothelial Carcinoma. The Journal of urology, 2026. 215(3).

37. K, Y., et al., A Spatial Atlas of Muscle-Invasive Bladder Cancer Reveals Lineage-Specific Vulnerabilities and Immune Architecture. Cancer discovery, 2026.

38. Rosenberg, J., et al., Atezolizumab monotherapy for metastatic urothelial carcinoma: final analysis from the phase II IMvigor210 trial - PubMed. ESMO open, 2024. 9(12).

39. Meylan, M., et al., Tertiary lymphoid structures generate and propagate anti-tumor antibody-producing plasma cells in renal cell cancer. Immunity, 2022. 55(3).

40. Rosenberg, J., et al., Atezolizumab monotherapy for metastatic urothelial carcinoma: final analysis from the phase II IMvigor210 trial. ESMO open, 2024. 9(12).

41. Liu, N., et al., standR: spatial transcriptomic analysis for GeoMx DSP data. Nucleic acids research, 2024. 52(1).

42. Tan, Y., et al., SPACEc: a streamlined, interactive Python workflow for multiplexed image processing and analysis. Nature Communications, 2025. 16(1).

43. Greenwald, N., et al., Whole-cell segmentation of tissue images with human-level performance using large-scale data annotation and deep learning. Nature biotechnology, 2021. 40(4).

44. Lee, M., et al., CellSeg: a robust, pre-trained nucleus segmentation and pixel quantification software for highly multiplexed fluorescence images. BMC bioinformatics, 2022. 23(1).

